# A frameshift mutation in *GON4L* is associated with proportionate dwarfism in Fleckvieh cattle

**DOI:** 10.1101/036889

**Authors:** Hermann Schwarzenbacher, Christine Wurmser, Krzysztof Flisikowski, Lubica Misurova, Simone Jung, Martin C. Langenmayer, Angelika Schnieke, Gabriela Knubben-Schweizer, Ruedi Fries, Hubert Pausch

## Abstract

**Background:** Low birth weight and postnatal growth restriction are the most evident symptoms of dwarfism. Accompanying skeletal aberrations may compromise the general condition and locomotion of affected individuals. Several paternal half sibs with low birth weight and size were born in 2013 in the Fleckvieh cattle population.

**Results:** Affected calves were strikingly underweight at birth despite a normal gestation length and had craniofacial abnormalities such as elongated narrow heads and brachygnathia inferior. Despite a normal general condition, their growth remained restricted during rearing. We genotyped 27 affected and 10,454 unaffected animals at 44,672 SNPs and performed association testing followed by homozygosity mapping to map the growth failure to a 1.85 Mb segment on bovine chromosome 3. Analysis of whole-genome re-sequencing data from one affected and 289 unaffected animals revealed a 1bp deletion (g.15079217delC, rs723240647) in the coding region of the GON4L gene that segregated with the dwarfism-associated haplotype. We show that the deletion introduces intron retention and premature termination of translation, putatively resulting in a severely truncated protein that lacks domains that are likely essential to normal protein function. The widespread use of an unnoticed carrier bull for artificial insemination has resulted in a tenfold increase in the frequency of the deleterious allele in the female population.

**Conclusions:** A frameshift mutation in *GON4L* is associated with autosomal recessive proportionate dwarfism in Fleckvieh cattle. The mutation has segregated in the population for more than fifty years without being recognized as a genetic disorder. However, the widespread use of an unnoticed carrier bull for artificial insemination caused a sudden accumulation of homozygous calves with dwarfism. Our findings provide the basis for genome-based mating strategies to avoid the inadvertent mating of carrier animals and thereby prevent the birth of homozygous calves with impaired growth.

## Background

Bovine stature is a prototypical complex trait that is controlled by a few loci with large effects and numerous loci with small effects. Genome-wide association studies using dense molecular markers revealed several quantitative trait loci (QTL) for growth-related traits in cattle [1-3]. The identified QTL account for a reasonable fraction of the phenotypic variation of bovine height [2, 4]. Sequence variants associated with mature height may also affect the size and weight of newborn calves [2, 3, 5].

Birth size and weight varies between breeds, parities and the sex of the calf [6, 7]. The birth weight in Fleckvieh cattle typically ranges from 38 to 45 kg [8]. Calves with strikingly low birth weight and size despite normal gestation length are commonly referred to as “dwarfs”.

Dwarfism (DW) has been observed in several cattle breeds including Fleckvieh [9-11]. Low birth size and postnatal growth restriction are the most apparent characteristics of DW. Undersized animals may be normally proportionated and have an undisturbed general condition (*i.e*., proportional DW [12]). However, DW may also be accompanied by disproportionately shortened limbs and skeletal deformities (*i.e*., disproportional DW, chondrodysplasia [13]). Depending on the severity of the structural aberrations, disproportionate DW may be fatal [14, 15].

Both autosomal recessive and dominant inheritance has been reported for bovine DW (*e.g*., [12, 15]). Causative mutations for DW were identified in Angus (OMIA 001485-9913 [16]), Dexter (OMIA 001271-9913 [14]), Tyrolean Grey (OMIA 000187-9913 [13]), Holstein-Friesian (OMIA 001926-9913 [15]) and Japanese Brown cattle (OMIA 000187-9913 [17]). However, mutations causing DW have not yet been identified in Fleckvieh cattle.

Here we present the phenotypic and genetic characterization of autosomal recessive DW in Fleckvieh cattle. The use of genome-wide association testing, autozygosity mapping and massive re-sequencing data enabled us to identify a frameshift mutation in *GON4L* that is likely causal for the growth failure.

## Methods

### Animal ethics statement

Two animals were hospitalized at the animal clinic of Ludwigs-Maximilians-Universität München. Another two animals were pathologically examined at the Institute for Veterinary Disease Control (IVDC) of Austrian Agency for Health and Food Safety. One hospitalized calf was euthanized because of recurrent tympania with no prospect of improvement and subsequently necropsied. Tissue samples were collected during necropsy. All affected animals result from inadvertent carrier matings that happened in Fleckvieh farms. No ethical approval was required for this study.

### Animals

Twenty-seven paternal half sibs (16 male, 11 female) with strikingly low birth weight and postnatal growth restriction were inspected by breeding consultants at the age of three weeks to 18 months. Ear tissue samples were collected by breeding consultants and DNA was prepared following standard DNA extraction protocols.

### Genotyping, quality control and haplotype inference

Twenty-seven affected animals were genotyped with the Illumina BovineSNP50 v2 BeadChip interrogating genotypes at 54,609 SNPs. The per-individual call-rate ranged from 98.96% to 99.60% with an average call-rate of 99.33%. In addition, genotypes of 10,454 unaffected Fleckvieh animals were available from genotyping with the Illumina BovineSNP50 v1 BeadChip and the Illumina BovineHD BeadChip [18, 19]. The genotype data of cases and controls were combined and SNPs that were present in both datasets were retained for further analyses. Following quality control (minor allele frequency above 0.5%, no deviation from the Hardy-Weinberg equilibrium (P>0.0001), per-SNP and per-individual call-rate higher than 95%), 10,481 animals (27 affected, 10,454 unaffected) and 44,672 SNPs remained for association testing. The *Beagle* software [20] was used to impute sporadically missing genotypes and to infer haplotypes.

### Haplotype-based association testing

A sliding window consisting of 25 contiguous SNPs (corresponding to an average haplotype length of 1.42±0.43 Mb) was shifted along the genome in steps of 2 SNPs. Within each sliding window, all haplotypes with a frequency above 0.5% (N=787,232) were tested for association with the affection status using Fisher exact tests of allelic association. Haplotypes with a P value less than 6.35 × 10^-8^ (5 *%* Bonferroni-corrected significance threshold) were considered as significantly associated.

### Generation of sequence data

Genomic DNA was prepared from a frozen semen sample of the supposed founder (DWhet) and from an ear tissue sample of one affected animal (DW_hom_) following standard DNA extraction protocols. Paired-end libraries were prepared using the paired-end TruSeq DNA sample preparation kit (Illumina) and sequenced using the HiSeq 2500 instrument (Illumina). The resulting reads were aligned to the University of Maryland reference sequence of the bovine genome (UMD3.1 [21]) using the *BWA* software tool [22]. Individual files in SAM format were converted into BAM format using *SAMtools* [23]. Duplicate reads were marked with the MarkDuplicates command of *Picard Tools* [24]. To help identify the causal mutation, we used sequence data from another 288 unaffected animals from nine cattle breeds (Gelbvieh, Nordic Finncattle, Fleckvieh, Holstein-Friesian, Brown-Swiss, Original Braunvieh, Original Simmental, Red-Holstein, Ayrshire) that had been generated previously [25, 26].

### Variant calling and imputation

SNPs, short insertions and deletions were genotyped in DWh_om_, DWhet and 288 control animals from nine cattle breeds simultaneously using the multi-sample approach implemented in *mpileup* of *SAMtools* along with *BCFtools* [23]. *Beagle* phasing and imputation (see above) was used to improve the primary genotype calling by *SAMtools*. The detection of structural variants was performed in DW_hom_, DW_he_t and 203 sequenced control animals that had an average genome fold coverage above 10x using the *Pindel* software package with default settings [27].

### Identification of candidate causal variants

To identify mutations compatible with recessive inheritance of DW, all polymorphic sites within the DW-associated region were filtered for variants that met three conditions: (i) DW_hom_ was homozygous for the alternate allele, (ii) DW_he_t was heterozygous and (iii) all control animals were homozygous for the reference allele. Candidate causal variants were annotated using the *Variant Effect Predictor* tool [28, 29]. Sequence variants of 1147 animals from Run4 of the 1000 bull genomes project [15] were analyzed to obtain the genotype distribution of candidate causal variants in diverse bovine populations.

### Manual re-annotation of the bovine *GON4L-gene*

A coding mutation in the *GON4L* coding sequence (rs723240647) gene was associated with DW. Since the annotation of the bovine genome may be flawed, we manually re-annotated the genomic structure of *GON4L* (ENSBTAG00000020356) based on the University of Maryland (UMD3.1) bovine genome sequence assembly [21] and the Dana-Farber Cancer Institute bovine gene index release 12.0 [30] using the *GenomeThreader* software tool [31]. The *GenomeThreader* output was viewed and edited using the *Apollo* sequence annotation editor [32].

### Validation of candidate causal variants

PCR primers were designed to scrutinise the rs723240647 polymorphism using Sanger sequencing (see Additional File 1). Genomic PCR products were sequenced using the BigDye^®^ Terminator v1.1 Cycle Sequencing Kit (Life Technologies) on the ABI 3130x1 Genetic Analyzer (Life Technologies). Genotypes for rs723240647 and rs715250609 were obtained in 3,882 and 1,851 Fleckvieh animals, respectively, using KASP^TM^ (LGC Genomics) genotyping assays (see Additional file 1).

### Clinical and patholgical examination of four animals with DW

Two calves with DW were pathologically examined at the Institute for Veterinary Disease Control (IVDC) of Austrian Agency for Health and Food Safety at the age of 101 and 143 days. Another two calves with DW were admitted to the animal clinic at the age of 57 and 93 days. Initial examination (including weighing) was performed upon admission. The younger calf suffered from recurrent tympania and was euthanized four days after admission because of no prospect of improvement and subsequently necropsied. Tissue samples were collected during necropsy. The older calf was hospitalized for 400 days. Weight records were collected once a week.

### RT-PCR

Total RNA from lymph nodes, thymus, lung, heart, pancreas, liver, kidney and spleen of the euthanized animal was extracted from tissue samples using Trizol (Invitrogen) according to the manufacturer’s protocol with some modifications. After *DNaseI* treatment (Ambion), RNA was quantified using a NanoDrop ND-1000 (PeqLab) spectrophotometer, and RNA integrity determined by RNA Nano6000 Labchip (Agilent Technologies). Complementary DNA (cDNA) was synthesized using the SuperScript IV transcriptase (Thermo Fisher Scientific). *GON4L* mRNA was examined by RT-PCR using primers 1F - GAGTCAAGCAGCTCAAACCC and 1R - AGCCAAGTCAGTTTCTCCATT, which hybridize to exons 20 and 21 and amplify a 348 bp product based on the mRNA reference sequence (NCBI accession number: XM_010802911) of the bovine *GON4L*. The shorter version of exon 21 was amplified using reverse primer 2R - CTCAGACTCACCCTCCTGACTC. RT-PCR was performed in 20 ml reaction volumes containing diluted first-strand cDNA equivalent to 50 ng input RNA. PCR products were loaded on 2% agarose gels.

## Results

### Phenotypic manifestation of dwarfism

Twenty-seven calves (16 males and 11 females) with strikingly low birth weight (˜15kg) and size despite normal gestation length were noticed among the descendants of an artificial insemination bull that was used for more than 290,000 inseminations. Four affected calves were clinically and pathologically examined. At age 61, 97, 101 and 143 days, they were underweight at 42, 79, 53 and 51 kg, respectively. The calves had multiple craniofacial aberrations (*i.e*., brachygnatia inferior, elongated narrow heads, structural deformities of the muzzle) and spinal distortions. Wrinkled skin, areas with excessive skin and a disproportionately large head became evident during rearing (Figure 1, see Additional file 2). Although the general condition, feed intake and locomotion of the animals were normal, their growth remained restricted. The average weight gain of an affected animal during a hospitalisation period of 400 days was only 450 g per day, *i.e*., less than half the weight gain of healthy Fleckvieh bulls (Figure 1h). The growth of the sire and all dams was normal. Since both sexes were affected and most dams had a common ancestor, we hypothesized an autosomal recessive mode of inheritance. Dominant inheritance of DW was unlikely because less then 1% of the progeny were affected.

**Figure 1.**
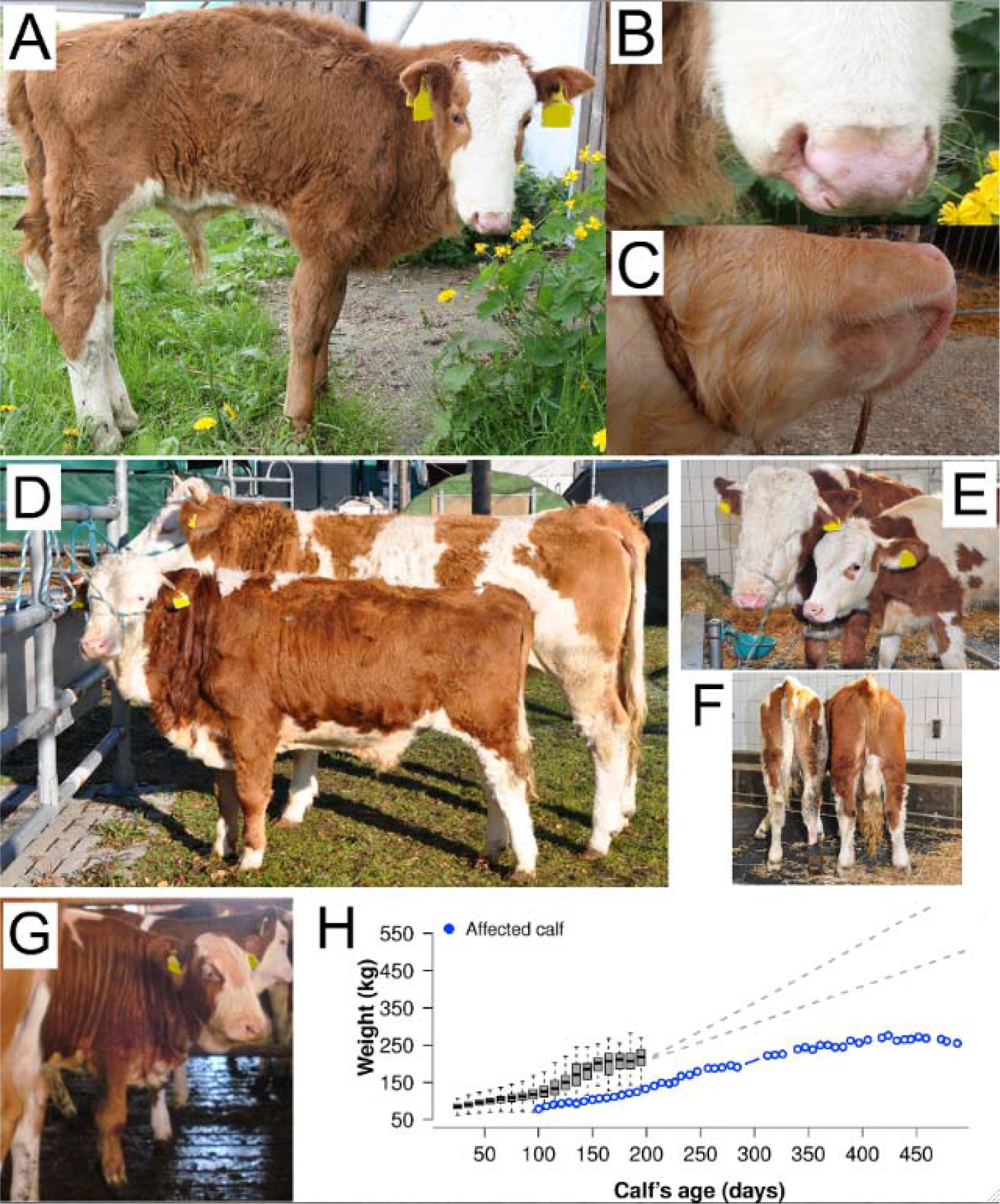
Phenotypic manifestation of dwarfism in Fleckvieh cattle. A twenty-week old Fleckvieh calf with crooked back, structural abnormalities of the muzzle and brachygnathia inferior (a-c). A fifteen-month old Fleckvieh bull with dwarfism and a healthy coeval (d). Note the skin flaps in the neck area. The same animal with a six-month old healthy animal (e, f). Note the disproportionately large head of the affected animal (left) (e) although its height (right animal, height at withers: 112 cm) is similar to the nine-month younger healthy animal (left) (f). An approximately 18-month old animal with dwarfism with excessive skin in the neck area (g). The weight of one animal with dwarfism (blue) is compared to the weight of 74,422 Fleckvieh calves (grey boxes) (h). The upper dotted line represents the growth of Fleckvieh bulls observed in Geuder et al. [46]. The lower dotted line is a growth curve assuming an average weight gain of 1000 g/day, *i.e*., a lower bound estimate for the growth of Fleckvieh bulls.

### Dwarfism maps to chromosome 3

To identify the genomic region associated with DW, 27 affected and 10,454 unaffected animals were genotyped using a medium-density genotyping array. After quality control, 44,672 SNPs were retained for genome-wide association testing. Because all affected animals were highly related with each other, the haplotype-based association study with DW revealed many significantly associated haplotypes. However, a striking association with DW of a proximal segment of bovine chromosome 3 became evident (Figure 2a). The most significant association signal (P=2.18 × 10^-124^) resulted from two contiguous haplotypes located between 14,884,969 bp and 16,557,950 bp.

**Figure 2.**
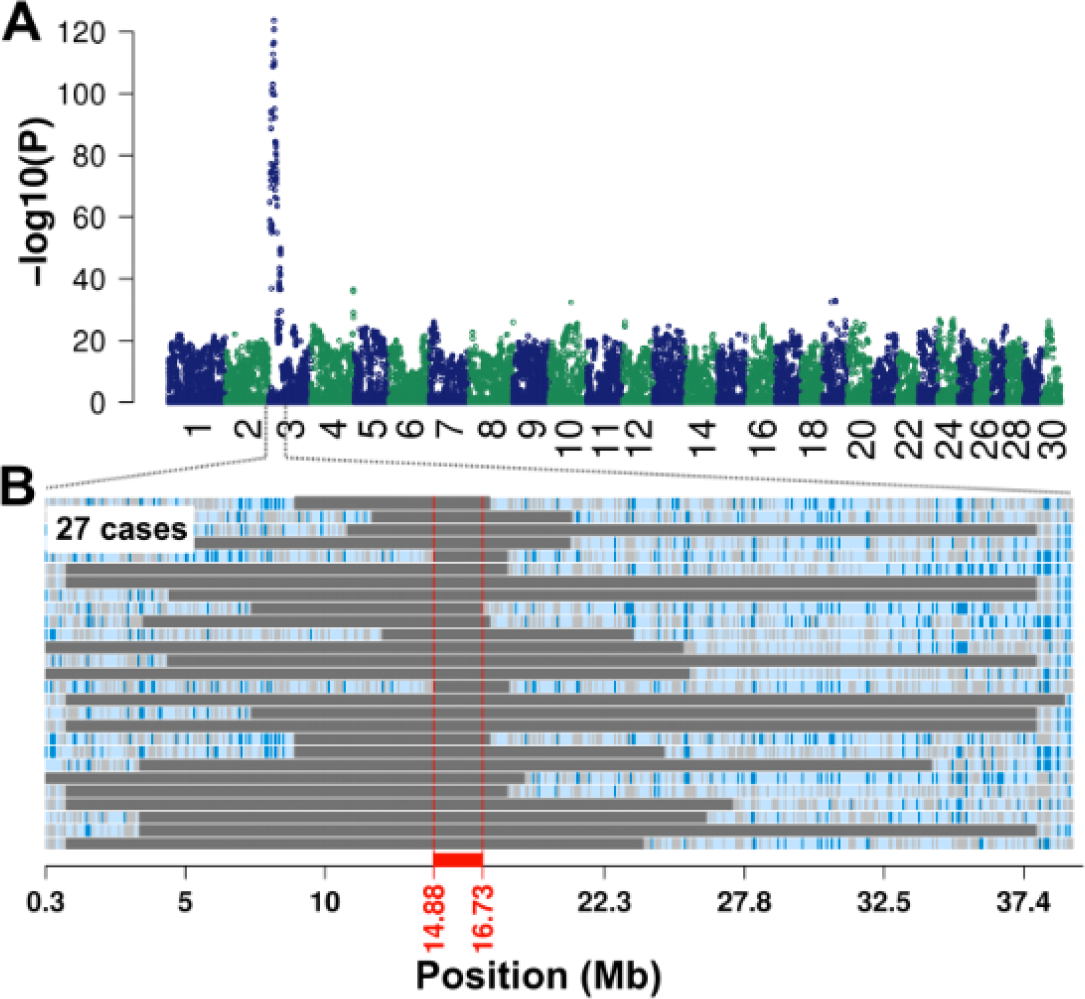
Mapping of the genomic region associated with dwarfism. Association of 787,232 haplotypes with dwarfism in 27 affected and 10,454 unaffected animals (a). P-values were obtained by calculating Fisher exact tests of allelic association. Autozygosity mapping in 27 animals with dwarfism (b). Blue and pale blue represent homozygous genotypes (AA and BB), heterozygous genotypes (AB) are displayed in light grey. The solid grey bars represent segments of extended homozygosity in 27 animals with dwarfism. The red bar indicates the common segment of homozygosity. Autozygosity mapping revealed a common 1.85 Mb segment (14.88 Mb - 16.73 Mb) of extended homozygosity in 27 affected animals, corroborating recessive inheritance (Figure 2b). The common segment of extended homozygosity encompassed 71 transcripts/genes. However, none of them was previously associated with DW.

Of 10,454 control animals, 81 were heterozygous and none was homozygous for the DW-associated haplotype, corresponding to a haplotype frequency of 0.38%. In the recent male breeding population (birth years 2000-2012), the frequency of the DW-associated haplotype was 0.25% (see Additional file 3). The haplotype frequency was considerably higher (2.6%) in the female population because of the widespread use of an unnoticed carrier bull for artificial insemination [33].

Haplotype and pedigree analysis enabled us to track the DW-associated haplotype back (up to twelve generations) to an artificial insemination bull (DW_he_t) born in 1959. DWhet was present in the maternal and paternal lineage of 21 affected animals. However, DW_he_t was not in the pedigree of six dams, possibly due to incomplete pedigree information and recording errors (see Additional file 4). The missing connection of six dams to DW_he_t may also indicate that the mutation occurred several generations before DW_het_.

### Identification of candidate causal variants for dwarfism

One affected animal (DW_hom_) and DW_het_ were sequenced to an average read depth of 13x. To help identify the underlying mutation, we additionally exploited sequence data of 288 animals from nine breeds including 149 Fleckvieh animals. None of 149 sequenced control animals of the Fleckvieh population carried the DW-associated haplotype.

Multi-sample variant calling within the 1.85 Mb segment of extended homozygosity revealed 11,475 single nucleotide and short insertion and deletion polymorphisms as well as 3,158 larger structural variants. These 14,633 polymorphic sites were filtered for variants that were compatible with recessive inheritance that is homozygous for the alternate allele in DW_hom_, heterozygous in DW_het_ and homozygous for the reference allele in 288 control animals (assuming the mutation is recessive and specific for the Fleckvieh breed). This approach revealed ten candidate causal variants for DW (Table 1). Five of them were intergenic, four were located in introns of *KCNN3, ADAR* and *TDRD10*, and one variant was located in the *GON4L* coding region (see Additional file 5).

Eight of ten compatible variants were excluded as being causative for DW because they segregated in 1005 animals from 28 breeds other than Fleckvieh that had been sequenced for the 1000 bull genomes project [15] (Table 1, see Additional file 6). In conclusion, only an intron variant in *TDRD10* (rs715250609) and a coding variant in *GON4L* (rs723240647) segregated with DW. The intron variant in *TDRD10* is unlikely to be deleterious to protein function because it is more than 4000 bp distant from the most proximal splice site. We therefore considered the coding variant in *GON4L* (gon-4-like) as the most likely causal mutation for DW.

**Table 1.**
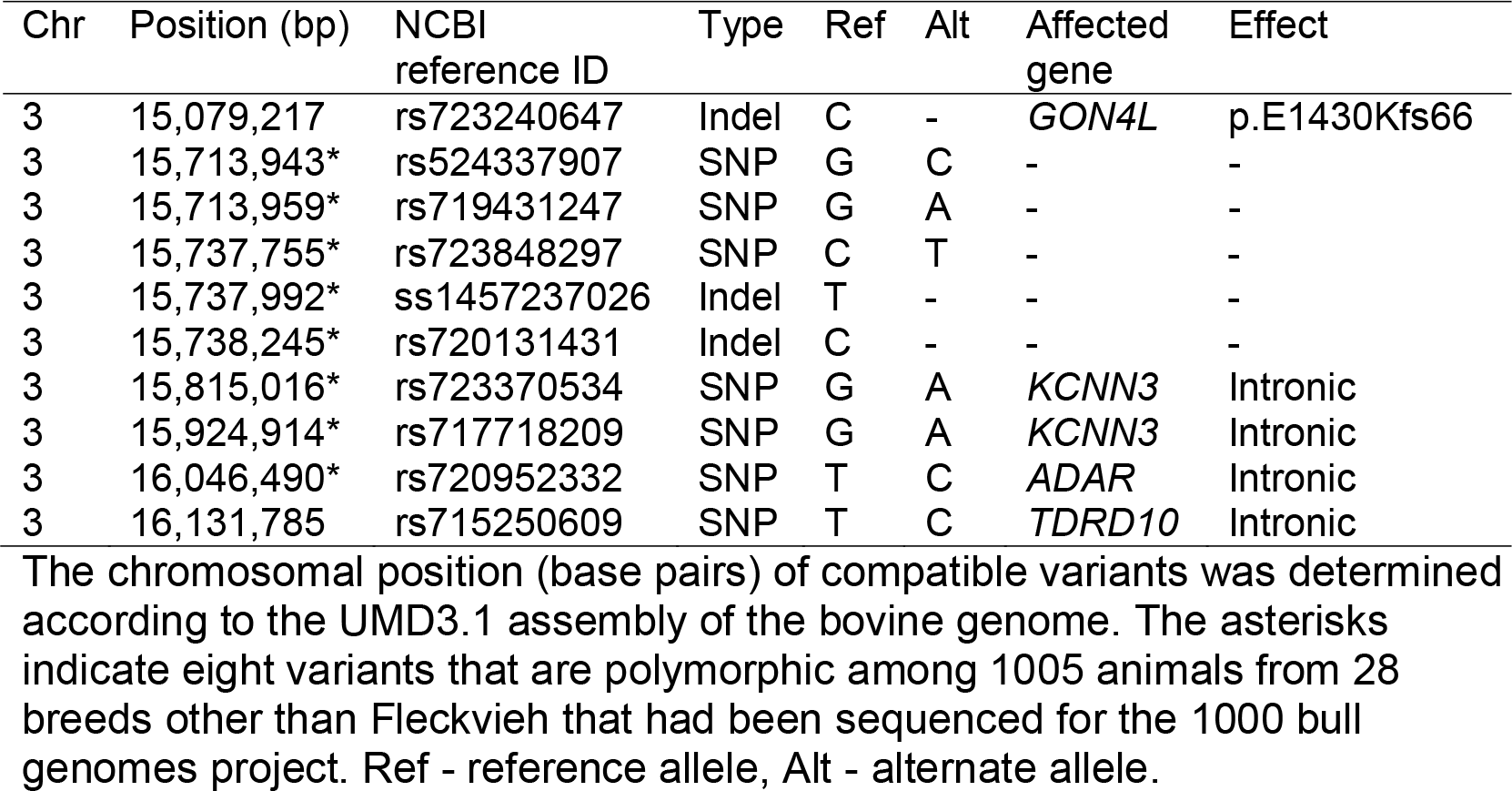
Ten sequence variants compatible with recessive inheritance. The chromosomal position (base pairs) of compatible variants was determined according to the UMD3.1 assembly of the bovine genome. The asterisks indicate eight variants that are polymorphic among 1005 animals from 28 breeds other than Fleckvieh that had been sequenced for the 1000 bull genomes project. Ref - reference allele, Alt - alternate allele.

| Chr | Position (bp) | NCBI<br>reference ID | Type | Ref | Alt | Affected<br>gene | Effect |
| --- | --- | --- | --- | --- | --- | --- | --- |
| 3 | 15,079,217 | rs723240647 | Indel | C | - | *GON4L* | p.E1430Kfs66 |
| 3 | 15,713,943* | rs524337907 | SNP | G | C | - | - |
| 3 | 15,713,959* | rs719431247 | SNP | G | A | - | - |
| 3 | 15,737,755* | rs723848297 | SNP | C | T | - | - |
| 3 | 15,737,992* | ss1457237026 | Indel | T | - | - | - |
| 3 | 15,738,245* | rs720131431 | Indel | C | - | - | - |
| 3 | 15,815,016* | rs723370534 | SNP | G | A | *KCNN3* | Intronic |
| 3 | 15,924,914* | rs717718209 | SNP | G | A | *KCNN3* | Intronic |
| 3 | 16,046,490* | rs720952332 | SNP | T | C | *ADAR* | Intronic |
| 3 | 16,131,785 | rs715250609 | SNP | T | C | *TDRD10* | Intronic |

### A 1bp deletion in *GON4L* is associated with dwarfism

Bovine *GON4L* consists of 31 exons encoding 2239 amino acids. The variant compatible with recessive inheritance is a 1bp deletion (rs723240647, g.15079217delC, ENSBTAT00000027126:c.4285_4287delCCCinsCC) in exon 20 (Figure 3a). Sanger sequencing confirmed that DWh_om_ and DW_he_t were homozygous and heterozygous, respectively, for g.15079217delC. The deletion introduces a frameshift in translation predicted to alter the protein sequence from amino acid position 1430 onwards, and a premature translation termination codon at position 1496 (p.Glu1430LysfsX66). The Gon-4-like protein contains highly conserved paired amphipathic helix (PAH) repeats and caspase 8-associated protein 2 myb-like (CASP8AP2) domains. The mutant protein is predicted to be shortened by 745 amino acids (33%) and lack domains that are likely essential for normal protein function (Figure 3b).

**Figure 3.**
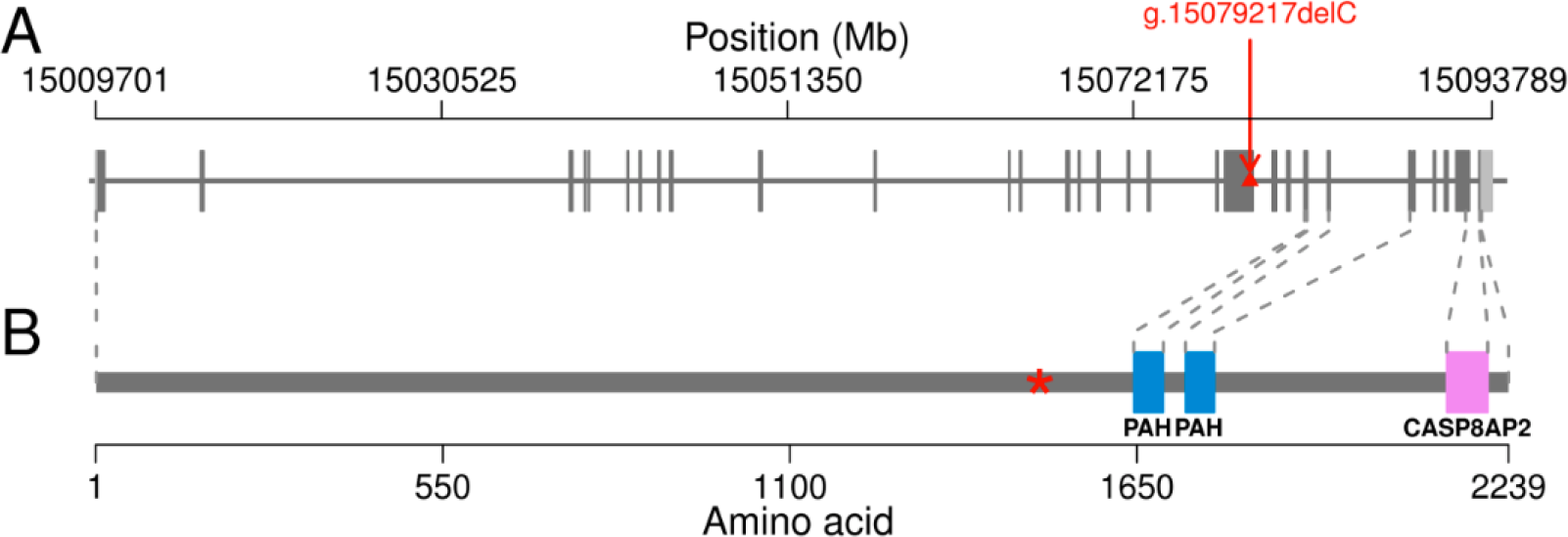
A 1bp deletion in *GON4L* introduces a premature stop codon. Genomic structure of the bovine gon-4-like encoding gene *GON4L* (a). Grey and light grey boxes represent exons and untranslated regions, respectively. The red triangle represents a 1bp deletion (rs723240647, g.15079217delC) in exon 20 of *GON4L*. Gon-4-like consists of 2239 amino acids and contains highly conserved paired amphipathic helix (PAH) repeat and caspase 8-associated protein 2 myb-like (CASP8AP2) domains (b). The red star indicates the premature stop codon resulting from the 1bp deletion.

Genotypes for rs723240647 and rs715250609 were obtained in cases and a large number of randomly selected unaffected Fleckvieh animals using customized KASP genotyping assays (Table 2). rs723240647 was highly significantly associated with DW (P=1.55 × 10). Twenty-seven calves with DW were homozygous for the deletion variant, while 3,855 unaffected animals were heterozygous and homozygous for the reference allele. One animal that carried the DW-associated haplotype was homozygous for the reference allele possibly indicating a swapped DNA sample, haplotype recombination or imperfect genotype phasing. The intron variant in *TDRD10* (rs715250609) was in almost complete linkage disequilibrium (r^2^=0.98) with rs723240647 (Table 2).

**Table 2:**
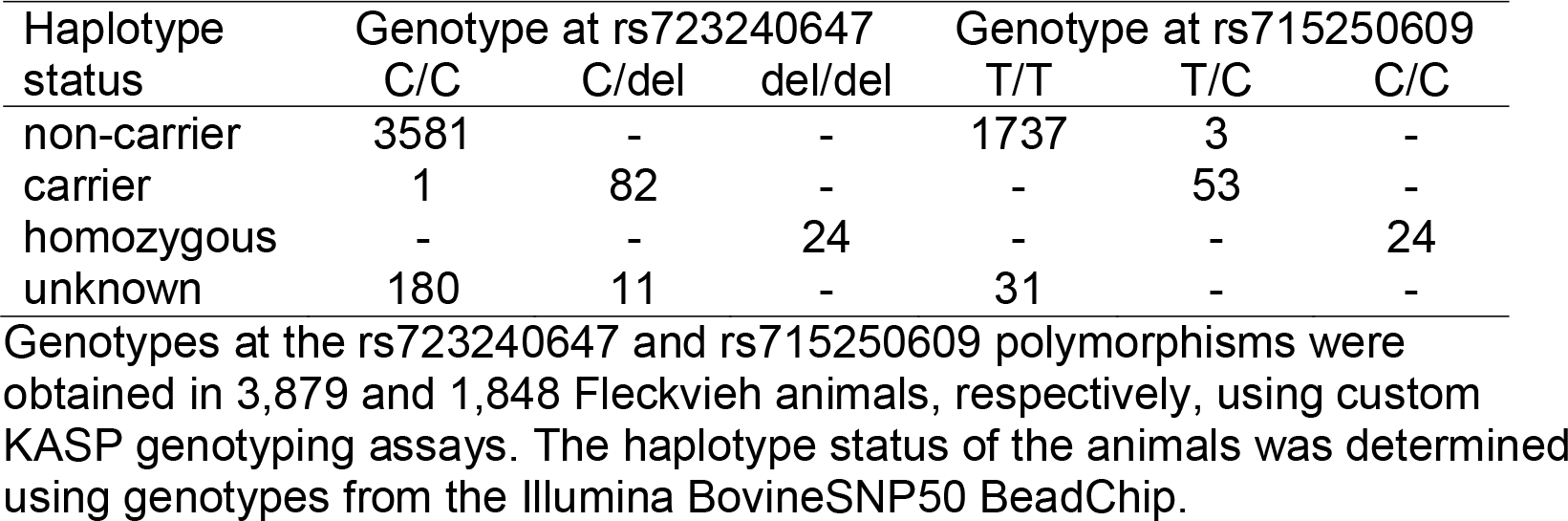
Genotypes for two mutations segregating with the DW-associated haplotype. Genotypes at the rs723240647 and rs715250609 polymorphisms were obtained in 3,882 and 1,851 Fleckvieh animals, respectively, using custom KASP genotyping assays. The haplotype status of the animals was determined using genotypes from the Illumina BovineSNP50 BeadChip.

### The deletion in *GON4L* creates intron retention and mRNA degradation

The effect of the g.15079217delC variant on *GON4L* transcription was examined by RT-PCR using RNA extracted from several tissues of a homozygous animal. Using primers located in exons 20 and 21, we obtained two RT-PCR products of 348 bp and ˜310 bp from a wild type and a mutant homozygous animal, respectively. The longer PCR fragment corresponded to the reference mRNA sequence (NM_001192626) of the bovine *GON4L* gene. The ˜310 bp PCR fragment showed a superimposed sequence of 35 bp at the 5’ end of exon 21, suggesting the presence of an alternative variant of exon 21, which is not directly associated with DW. The presence of different isoforms in the 3’ terminal end of *GON4L* in human and cattle have been reported previously. The intensity of the alternative cDNA fragment was higher in the mutant homozygote than in the wild type animal, possibly indicating degradation of the mutant transcript in the homozygous animal (see Additional file 7). We designed a reverse RT-PCR primer specific for the alternative exon 21, and obtained a unique 348 bp RT-PCR product from the wild type animal and two RT-PCR products of 313 bp and ˜1500 bp from the mutant homozygous animal (Figure 4). DNA sequence analysis of the 348 bp wild type RT-PCR product revealed that it corresponded to the mRNA reference sequence of the bovine *GON4L* gene. Sequence analysis of the longer fragment in the mutant homozygous animal revealed retention of intron 20. The length of the longer PCR fragment was 1488 bp. The retention of intron 20 is predicted to introduce a frameshift resulting in a premature translation termination codon at position 1492. In conclusion, the animal homozygous for the g.15079217delC variant contains the premature translation termination codon at position 1492.

**Figure 4.**
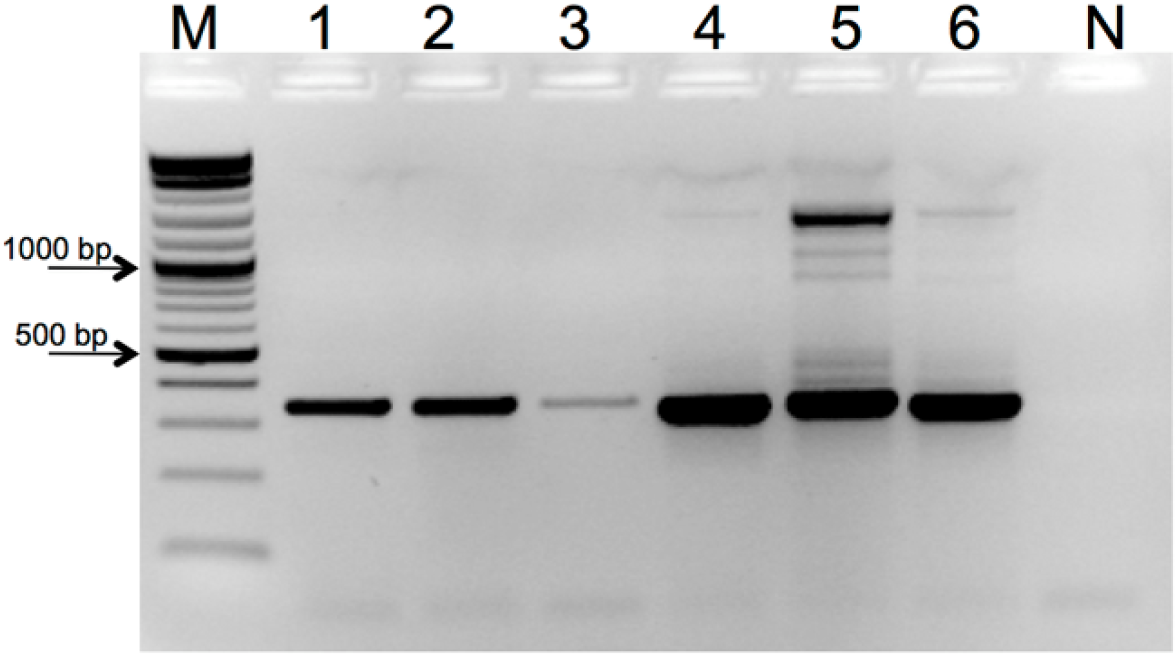
Retention of intron 20 of the *GON4L* gene in a Fleckvieh calf with dwarfism. (a) RT-PCR products were separated by 2% agarose gel electrophoresis. Lanes 1, 2, and 3 show 348 bp RT-PCR products from lung, lymph nodes, and liver of a wild type animal. Lines 4, 5, and 6 show 348 bp and 1488 bp RT-PCR products from lung, lymph nodes, and liver of a homozygous calf. M - size marker, N - negative RT-PCR control. Schematic representation of exons 19 to 22 of wild type (b) and mutant (c) *GON4L*. Blue arrows represent forward (1F) and reverse primers (1R, 2R). The red ‘X’ displays the premature translation termination at amino acid position 1492 resulting from the retention of intron 20. The light grey colour (c) displays intron 20.

## Discussion

A 1bp deletion in *GON4L* (g.15079217delC) was associated with DW in Fleckvieh cattle. The g.15079217delC variant introduces intron retention and premature translation termination resulting in a truncated protein. Compared to the wild type variant, the mutant GON4L protein is shortened by more than 30%. RNA analysis indicated that the mutant protein variant is less abundant, suggesting that it may be degraded *via* nonsense-mediated mRNA decay. If the truncated protein is (partially) retained, however, its function may be compromised because it lacks domains that are possibly essential for normal protein function. Loss-of-function variants in *Udu*, a gene that is highly homologous to *GON4L*, compromise cell cycle progression and response to DNA damage and thereby disturb embryonic growth in *D. rerio* [34-37]. In our study, the g.15079217delC variant was also associated with prenatal growth failure as evidenced by strikingly low birth weight of homozygous calves. The phenotypic manifestation of homozygosity for g.15079217delC, *i.e*., pre-and postnatal growth restriction and craniofacial aberrations, resembles phenotypic patterns of human primordial DW resulting from DNA repair disorders [38, 39]. Such findings suggest that disturbed growth of homozygous animals might result from defective responses to DNA damage due to impaired GON4L function. However, the actual mechanism(s) and pathway(s) that cause the extremely low birth weight and postnatal growth restriction of homozygous animals have yet to be elucidated.

Congenital disorders that manifest as growth failure have been identified in several cattle breeds. Affected calves may be born underweight or fail to thrive during rearing [26, 40-42]. Homozygosity for g.15079217delC becomes evident at birth. Unlike mutations in *ACAN* and *COL2A1* that cause lethal disproportionate DW in cattle [14, 15], homozygosity for g.15079217delC is not fatal. Apart from large heads, affected animals were normally proportionate. Moreover, their general condition and locomotion was normal and their weight gain was constant, although considerably less than healthy animals. Thus, homozygosity for g.15079217delC is less detrimental than, *e.g*., homozygosity for a mutation in *EVC2* which compromises both growth and locomotion of affected animals [13]. Nevertheless, homozygous animals are more likely to be culled at juvenile ages because of their reduced growth performance.

The g.15079217delC variant has segregated in the Fleckvieh population for more than fifty years, but due to its low frequency DW was rarely reported. Assuming a frequency of 0.2% of the deleterious allele, equal use of all bulls and 1,500,000 annual births in the German and Austrian Fleckvieh populations, one would expect only six homozygous calves with DW per year. However, the widespread use of unnoticed carriers of rare recessive alleles in artificial insemination may cause a sudden accumulation of affected calves, as our study demonstrates. Twenty-seven calves with DW were descendants from a bull that was used for more than 290,000 inseminations. The heavy use of this carrier bull resulted in a more than tenfold increase in allele frequency in the female population [33]. Our findings now enable the rapid identification of carrier animals. The g.15079217delC variant was in almost complete linkage disequilibrium with the DW-associated haplotype. Only one animal was misclassified using haplotype information demonstrating a high sensitivity and specificity of haplotype-based identification of DW-mutation carriers. Since all male breeding animals are routinely genotyped with dense genotyping arrays, carriers can be readily identified using haplotype information. However, only direct gene tests will unequivocally distinguish between carrier and non-carrier animals [43]. The identification of the frameshift mutation in *GON4L* now also enables the development of customized genotyping assays to identify carrier animals. Excluding carrier bulls from artificial insemination will prevent the emergence of homozygous animals and remove from the Fleckvieh population the rare DW-associated allele within a few generations. However, sophisticated strategies are required to simultaneously consider multiple deleterious alleles in genomic breeding programs while maintaining genetic diversity and high rates of genetic gain [44, 45].

## Conclusions

A frameshift variant in *GON4L* was associated with autosomal recessive proportionate DW in Fleckvieh cattle. The deleterious allele persisted in the Fleckvieh population for more than fifty years at very low frequency without being recognized as a genetic disorder. However, the heavy use of an unnoticed carrier bull for artificial insemination resulted in an accumulation of homozygous calves with DW and a tenfold increase in frequency of the deleterious allele in the female population. Our results provide the basis for the rapid identification of carrier bulls and the implementation of genome-based mating strategies to avoid inadvertent carrier matings, thereby preventing the birth of homozygous calves with unsatisfactory growth performance.

## Availability of supporting data

Whole-genome sequencing data of DWhom and DWhet were deposited in the European Nucleotide Archive (http://www.ebi.ac.uk/ena) under accession number PRJEB12832.

## Authors’ contribution

HP, and HS conceived the study; HP analyzed pedigree, genotyping and sequencing data; HS analyzed genotyping data and sampled affected animals; LM, MCL and GKS carried out clinical and pathological examinations; CW carried out the sequencing experiments; KF, SJ, ASch carried out the molecular genetic experiments; RF analyzed sequencing data; HP wrote the manuscript with the input from all authors. All authors read and approved the manuscript.

## Acknowledgements

We acknowledge Arbeitsgemeinschaft Süddeutscher Rinderzüchter und Besamungsorganisationen e.V. (ASR), Arbeitsgemeinschaft österreichischer Fleckviehzüchter (AGÖF) and Förderverein Bioökonomieforschung e.V. (FBF) for providing genotyping data. We acknowledge Dr. Josef Miesenberger from Oberösterreichische Besamungsstation GmbH for support in sample collection and Guido Däumler from Office for Food Agriculture and Forestry Ansbach for providing photographs of affected calves. We thank Alexander Kind for help with text editing of the manuscript. This work was supported by the German Research Foundation (DFG) and the Technische Universität München within the funding programme Open Access Publishing.

## Additional Files

Additional file 1

Title: Primer sequences used for the validation of two candidate causal mutations.

Additional file 2

Title: Fleckvieh calves with dwarfism.

Description: Affected calves with crooked back (a, b), elongated narrow heads and brachygnathia inferior (c-e). Wrinkled skin and areas with excessive skin (particularly in the neck area) became evident during rearing (c, e, g). A 19-month old animal with dwarfism and an eleven-month old healthy animal (f). The head of the affected animal was disproportionately large compared to its body.

Additional file 3

Title: Frequency of the dwarfism-associated haplotype in 8332 Fleckvieh bulls. Description: Grey bars and black dots represent the number of genotyped bulls and the haplotype frequency, respectively, as a function of the birth year.

Additional file 4

Title: Analysis of pedigree records of 27 animals with dwarfism.

Description: Pedigree of 21 animals with dwarfism (a). The pedigree contains only obligate mutation carriers. Red and yellow colour indicates 21 affected animals and 14 genotyped haplotype carriers, respectively. Green and blue indicates the sire of 21 affected animals and DWhet, respectively. Rectangles and ovals represent male and female animals, respectively. Pedigree completeness of another six animals with dwarfism with no link to the bull DWhet on the maternal path (b).

Additional file 5

Title: Annotation of ten candidate causal variants.

Description: The functional consequence of ten candidate causal variants was obtained from Ensembl using the *Variant Effect Predictor* tool.

Additional file 6

Title: Genotype distribution of ten candidate causal mutations for dwarfism in 1147 animals from the 1000 bull genomes project.

Description: Alternate allele frequency and genotype distribution of ten variants in 29 breeds (homozygous animals for the reference allele | heterozygous animals | homozygous animals for the alternate allele). Blue color indicates two variants that were not polymorphic among 1147 sequenced animals.

Additional file 7

Title: Electrophoregram presenting the *GON4L* cDNA sequencing result with superimposed sequence at the 5’ terminal end of exon 21 (marked in box).

## References

1. Karim L, Takeda H, Lin L, Druet T, Arias JAC, Baurain D, at al. Variants modulating the expression of a chromosome domain encompassing PLAG1 influence bovine stature. Nat Genet. 2011; 43: 405–13.

2. Pausch H, Flisikowski K, Jung S, Emmerling R, Edel C, Götz K-U, et al. Genome-wide association study identifies two major loci affecting calving ease and growth-related traits in cattle. Genetics. 2011; 187: 289–97.

3. Utsunomiya YT, Carmo AS do, Carvalheiro R, Neves HH, Matos MC, Zavarez LB, et al. Genome-wide association study for birth weight in Nellore cattle points to previously described orthologous genes affecting human and bovine height. BMC Genetics. 2013; 14: 52.

4. Saatchi M, Schnabel RD, Taylor JF, Garrick DJ. Large-effect pleiotropic or closely linked QTL segregate within and across ten US cattle breeds. BMC Genomics. 2014; 15: 442.

5. Riley DG, Welsh TH, Gill CA, Hulsman LL, Herring AD, Riggs PK, et al. Whole genome association of SNP with newborn calf cannon bone length. Livestock Sci. 2013; 155: 186–96.

6. Comerford JW, Bertrand JK, Benyshek LL, Johnson MH. Reproductive rates, birth weight, calving ease and 24-h calf survival in a four-breed diallel among Simmental, Limousin, Polled Hereford and Brahman beef cattle. J Anim Sci. 1987; 64: 65–76.

7. Johanson JM, Berger PJ. Birth weight as a predictor of calving ease and perinatal mortality in Holstein cattle. J Dairy Sci. 2003; 86: 3745–55.

8. Brandt H, Müllenhoff A, Lambertz C, Erhardt G, Gauly M. Estimation of genetic and crossbreeding parameters for preweaning traits in German Angus and Simmental beef cattle and the reciprocal crosses. J Anim Sci. 2010; 88: 80–6.

9. Johnson LE, Harshfield GS, McCONE W. Dwarfism, a hereditary defect in beef cattle. J Hered. 1950, 41: 177–81.

10. Becker RB, Neal FC, Wilcox CJ. Prenatal achondroplasia in a Jersey. J Dairy Sci. 1969; 52: 1122–3.

11. Gottwald VW. Über das Vorkommen von Zwergwuchs in der Nachzucht eines Fleckviehbullen. Reprod Domest Anim. 1967; 2: 63–7.

12. Latter MR, Latter BDH, Wilkins JF, Windsor PA. Inheritance of proportionate dwarfism in Angus cattle. Aust Vet J.. 2006; 84: 122–8.

13. Murgiano L, Jagannathan V, Benazzi C, Bolcato M, Brunetti B, Muscatello LV, et al. Deletion in the EVC2 gene causes chondrodysplastic dwarfism in Tyrolean Grey cattle. PLoS ONE. 2014; 9: e94861.

14. Cavanagh JAL, Tammen I, Windsor PA, Bateman JF, Savarirayan R, Nicholas FW, et al. Bulldog dwarfism in Dexter cattle is caused by mutations in ACAN. Mamm Genome. 2007; 18: 808–14.

15. Daetwyler HD, Capitan A, Pausch H, Stothard P, van Binsbergen R, Brøndum RF, et al. Whole-genome sequencing of 234 bulls facilitates mapping of monogenic and complex traits in cattle. Nat Genet. 2014; 46: 858–65.

16. Koltes JE, Mishra BP, Kumar D, Kataria RS, Totir LR, Fernando RL, et al. A nonsense mutation in cGMP-dependent type II protein kinase (PRKG2) causes dwarfism in American Angus cattle. Proc Natl Acad Sci USA.. 2009; 106: 19250–5.

17. Takeda H, Takami M, Oguni T, Tsuji T, Yoneda K, Sato H, et al. Positional cloning of the gene LIMBIN responsible for bovine chondrodysplastic dwarfism. P Natl Acad Sci USA. 2002; 99: 10549–54.

18. Pausch H, Kölle S, Wurmser C, Schwarzenbacher H, Emmerling R, Jansen S, et al. A nonsense mutation in TMEM95 encoding a nondescript transmembrane protein causes idiopathic male subfertility in cattle. PLOS Genetics. 2014; 10: e1004044.

19. Ertl J, Edel C, Emmerling R, Pausch H, Fries R, Götz K-U. On the limited increase in validation reliability using high-density genotypes in genomic best linear unbiased prediction: Observations from Fleckvieh cattle. J Dairy Sci. 2014; 97: 487–96.

20. Browning BL, Browning SR. A unified approach to genotype imputation and haplotype-phase inference for large data sets of trios and unrelated individuals. Am J Hum Genet. 2009; 84: 210–23.

21. Zimin AV, Delcher AL, Florea L, Kelley DR, Schatz MC, Puiu D, et al. A whole-genome assembly of the domestic cow, Bos taurus. Genome Biol. 2009; 10: R42–R42.

22. Li H, Durbin R. Fast and accurate short read alignment with Burrows-Wheeler transform. Bioinformatics. 2009; 25: 1754–60.

23. Li H, Handsaker B, Wysoker A, Fennell T, Ruan J, Homer N, et al The Sequence Alignment/Map format and SAMtools. Bioinformatic.s. 2009; 25: 2078–9.

24. Picard Tools - By Broad Institute. http://broadinstitute.github.io/picard/. Accessed 15 January 2016

25. Jansen S, Aigner B, Pausch H, Wysocki M, Eck S, Benet-Pagès A, et al. Assessment of the genomic variation in a cattle population by re-sequencing of key animals at low to medium coverage. BMC Genomics. 2013; 14: 446.

26. Pausch H, Schwarzenbacher H, Burgstaller J, Flisikowski K, Wurmser C, Jansen S, Homozygous haplotype deficiency reveals deleterious mutations compromising reproductive and rearing success in cattle. BMC Genomics. 2015; 16: 312.

27. Ye K, Schulz MH, Long Q, Apweiler R, Ning Z. Pindel: a pattern growth approach to detect break points of large deletions and medium sized insertions from paired-end short reads. Bioinformatics. 2009; 25: 2865–71.

28. McLaren W, Pritchard B, Rios D, Chen Y, Flicek P, Cunningham F. Deriving the consequences of genomic variants with the Ensembl API and SNP Effect Predictor. Bioinformatics. 2010; 26: 2069–70.

29. Variant Effect Predictor. http://www.ensembl.org/Tools/VEP. Accessed 15 January 2016

30. Quackenbush J, Cho J, Lee D, Liang F, Holt I, Karamycheva S, et al. The TIGR Gene Indices: analysis of gene transcript sequences in highly sampled eukaryotic species. Nucleic Acids Res. 2001; 29: 159–64.

31. Gremme G, Brendel V, Sparks ME, Kurtz S. Engineering a software tool for gene structure prediction in higher organisms. Inform Software Tech. 2005; 47: 965–78.

32. Lewis SE, Searle SMJ, Harris N, Gibson M, Lyer V, Richter J, et al. Apollo: a sequence annotation editor. Genome Biol. 2002; 3: 82.

33. Egger-Danner C, Schwarzenbacher H, Fuerst C, Willam A. Management of genetic disorders in the breeding program Fleckvieh AUSTRIA: results of model calculations. Zuechtungskunde. 2015; 87: 201–14.

34. Hammerschmidt M, Pelegri F, Mullins MC, Kane DA, Brand M, van Eeden FJ, et al. Mutations affecting morphogenesis during gastrulation and tail formation in the zebrafish, Danio rerio. Development. 1996; 123: 143–51.

35. Liu Y, Du L, Osato M, Teo EH, Qian F, Jin H, et al. The zebrafish udu gene encodes a novel nuclear factor and is essential for primitive erythroid cell development. Blood. 2007; 110: 99–106.

36. Lim C-H, Chong S-W, Jiang Y-J. Udu deficiency activates DNA damage checkpoint. Mol Biol Cell. 2009; 20: 4183–93.

37. Klingseisen A, Jackson AP. Mechanisms and pathways of growth failure in primordial dwarfism. Genes Dev. 2011; 25: 2011–24.

38. Woods CG. DNA repair disorders. Arch Dis Child. 1998; 78: 178–84.

39. Harley ME, Murina O, Leitch A, Higgs MR, Bicknell LS, Yigit G, et al. TRAIP promotes DNA damage response during genome replication and is mutated in primordial dwarfism. Nat Genet. 2016; 48: 36–43.

40. Hirano T, Kobayashi N, Matsuhashi T, Watanabe D, Watanabe T, Takasuga A, et al Mapping and exome sequencing identifies a mutation in the IARS gene as the cause of hereditary perinatal weak calf syndrome. PLoS ONE. 2013; 8: e64036.

41. Sartelet A, Druet T, Michaux C, Fasquelle C, Géron S, Tamma N, et al. A splice site variant in the bovine RNF11 gene compromises growth and regulation of the inflammatory response. PLoS Genet. 2012; 8: e1002581.

42. Jung S, Pausch H, Langenmayer MC, Schwarzenbacher H, Majzoub-Altweck M, Gollnick NS, et al. A nonsense mutation in PLD4 is associated with a zinc deficiency-like syndrome in Fleckvieh cattle. BMC Genomics. 2014; 15: 623.

43. Biffani S, Dimauro C, Macciotta N, Rossoni A, Stella A, Biscarini F. Predicting haplotype carriers from SNP genotypes in Bos taurus through linear discriminant analysis. Genet Sel Evol. 2015; 47: 4.

44. Segelke D, Täubert H, Reinhardt F, Thaller G. Considering genetic characteristics in German Holstein breeding programs. J Dairy Sci. 2016; 99: 458–67.

45. Cole JB: A simple strategy for managing many recessive disorders in a dairy cattle breeding program. Genet Sel Evol. 2015; 47: 94.

46. Geuder U, Pickl M, Scheidler M, Schuster M, Götz K. Mast-, Schlachtleistung und Fleischqualität bayerischer Rinderrassen. Zuechtungskunde. 2012; 36: 485–99.

